# PocketGNN: A Cross-Modal Framework Unifying Local 3D Pocket Geometry and Global Sequence Semantics for Enzyme Kinetic Prediction

**DOI:** 10.64898/2026.01.19.700244

**Authors:** Zihao Li, Diannan Lu

## Abstract

The enzyme turnover number (*k*_cat_) is a pivotal kinetic parameter for understanding bio-catalytic efficiency, yet its accurate prediction remains a grand challenge due to the complex interplay between local physicochemical constraints and global evolutionary context. Existing methods typically bifurcate into sequence-based approaches, which capture evolutionary semantics but miss fine-grained spatial details, or structure-based models, which often suffer from noise in whole-protein representations or lack global context.

To bridge this gap, we propose **PocketGNN**, a *cross-modal* deep learning framework that synergizes the precision of local 3D geometry with the breadth of global 1D sequence semantics. PocketGNN introduces a high-fidelity graph representation of the active pocket, enriched with a novel 24-dimensional geometric edge encoding (RBF distances, bond angles, dihedral angles) to capture the stereochemical determinants of catalysis. Crucially, this local structural view is fused with global evolutionary information extracted from pre-trained protein language models (ESM-2), creating a unified representation that spans spatial scales.

Evaluated on a rigorous dataset derived from IntEnzyDB, PocketGNN achieves a Pearson correlation coefficient (*r*) of **0.98** and a coefficient of determination (*R*^2^) of **0.918** for log_10_(*k*_cat_) under standard random splitting. Furthermore, under a strict 40% sequence identity split designed to test zero-shot generalization to unseen families, the model maintains a robust correlation (*r* = 0.67, *R*^2^ = 0.44), significantly outperforming recent state-of-the-art methods including CatPred (*r* = 0.52) and CataPro (*r* = 0.50). Interpretability analysis confirms that the model successfully attends to key catalytic residues, validating its ability to learn chemically meaningful structure-function relationships rather than mere sequence memorization.

## 1 Introduction

Enzymes are the orchestrators of life, driving metabolic flux with exquisite specificity. The catalytic turnover number (*k*_cat_) serves as the quantitative benchmark of this efficiency [1]. In the era of synthetic biology, the ability to predict *k*_cat_ in silico is a holy grail for rational enzyme design and metabolic pathway engineering [3, 11]. However, *k*_cat_ is an emergent property governed by factors across multiple scales: from the sub-angstrom precision of transition state stabilization in the active site to the global protein dynamics and stability encoded in the entire sequence.

### Related Work

Predicting enzyme kinetic parameters from sequence and structural information is challenging due to sparse measurements, heterogeneous assay conditions, and substantial experimental noise. Early deep-learning efforts such as **DLKcat** [2] combined protein sequence features with small-molecule representations to predict *k*_cat_ at scale, demonstrating that learned representations can capture enzyme–substrate-dependent trends. More recent works increasingly emphasize robust evaluation and broader kinetic coverage: **UniKP** [3] proposed a unified framework leveraging pretrained protein language models (pLMs) to predict multiple kinetic parameters. In parallel, **CatPred** [4] and **CataPro** [5] introduced comprehensive frameworks incorporating uncertainty estimates and broader parameterization, illustrating the field-wide trend toward multimodal feature integration.

A major driver of progress has been the emergence of large-scale protein language models like **ProtTrans** [6] and **ESM-2/ESMFold** [7]. These models demonstrate evolutionary-scale modeling and strong representational power, motivating the use of frozen pLM embeddings as global context features. Beyond sequence-only models, structure-aware architectures like **Uni-Mol** [8] and **PiFold** [9] have leveraged graph-based encoders for molecular representation and inverse folding. Recent advances such as **EZSpecificity** [10] employ cross-attention–empowered SE(3)-equivariant GNNs, demonstrating that explicitly modeling interactions between modalities can improve performance.

### Our Solution: PocketGNN

These developments suggest a converging direction: combining local 3D pocket geometry with global sequence-level evolutionary context. Therefore, we propose **PocketGNN**, a cross-modal framework designed to unify these complementary views. We posit that accurate kinetic prediction requires a “dual-lens” approach:

1. **Local High-Fidelity Geometry**: We model the active pocket not just as a bag of atoms, but as a geometrically constrained system. By incorporating bond angles and dihedral angles into the graph message passing, we explicitly capture the “short-range” physical constraints essential for catalysis.
2. **Global Evolutionary Semantics**: We acknowledge that the pocket does not exist in isolation. By fusing embeddings from ESM-2, we inject “long-range” evolutionary context (e.g., EC family priors, conservation patterns) into the prediction.

This fusion of *local 3D physics* and *global 1D evolution* allows PocketGNN to outperform existing baselines, providing a robust tool for functional enzyme mining.

## 2 Methods

### 2.1 Data Curation and Pipeline

We curate our dataset from IntEnzyDB [11], filtering for entries with valid UniProt IDs, substrate SMILES, and experimental *k*_cat_ values. To ensure data quality, we exclude extreme outliers and select entries where *k*_cat_ ∈ [10^−2^, 10^6^] s^−1^. Our final dataset comprises approximately 9,000 enzyme-substrate pairs with experimentally measured *k*_cat_ values, covering 6 major EC classes (EC 1-6).

The data processing pipeline (Figure 1) proceeds as follows:

**Figure 1.**
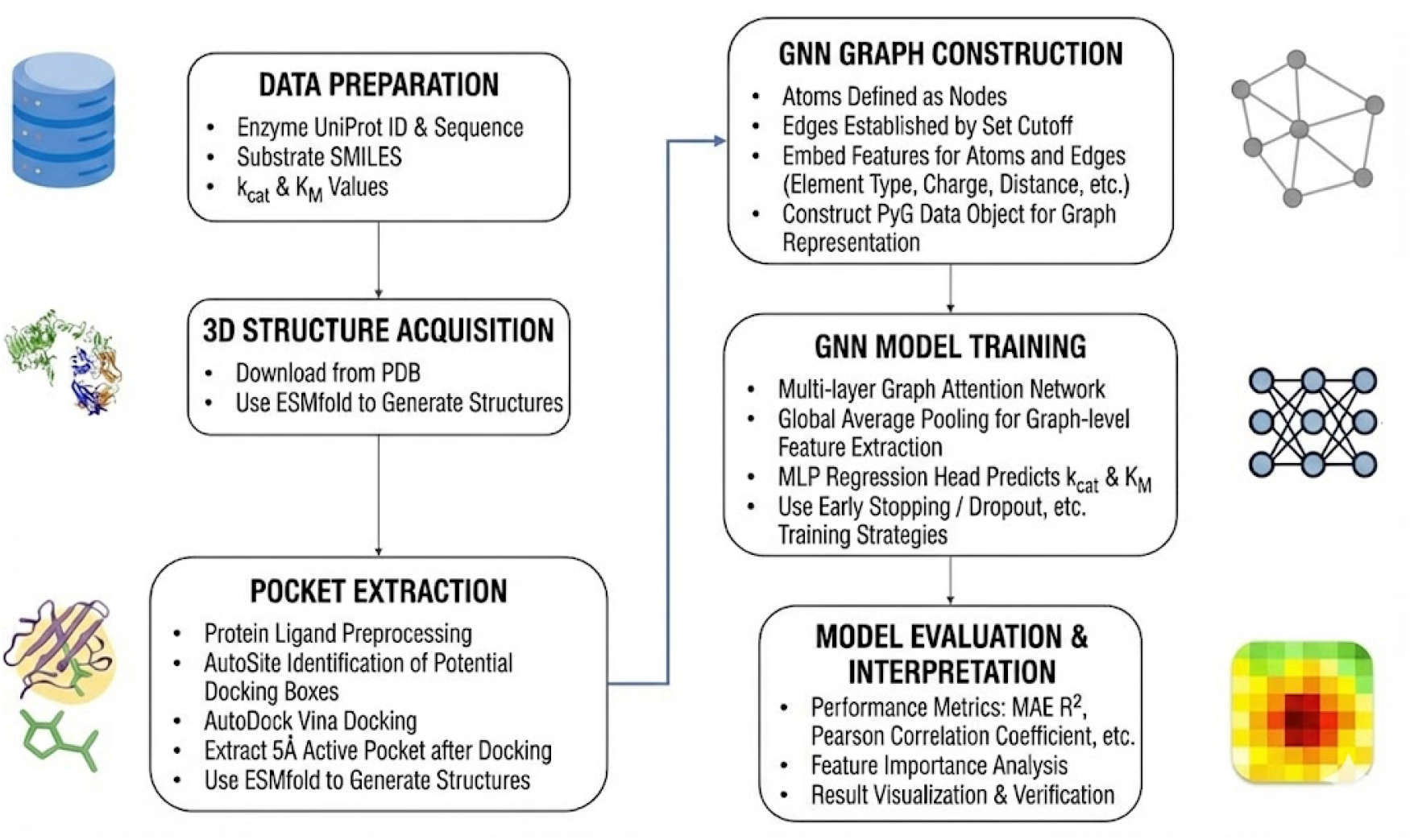
The cross-modal framework of PocketGNN. The model integrates a high-fidelity local graph encoder (capturing distances, angles, and torsions) with a global sequence encoder (ESM-2), bridging the gap between fine-grained geometry and evolutionary semantics.

**Figure 2.**
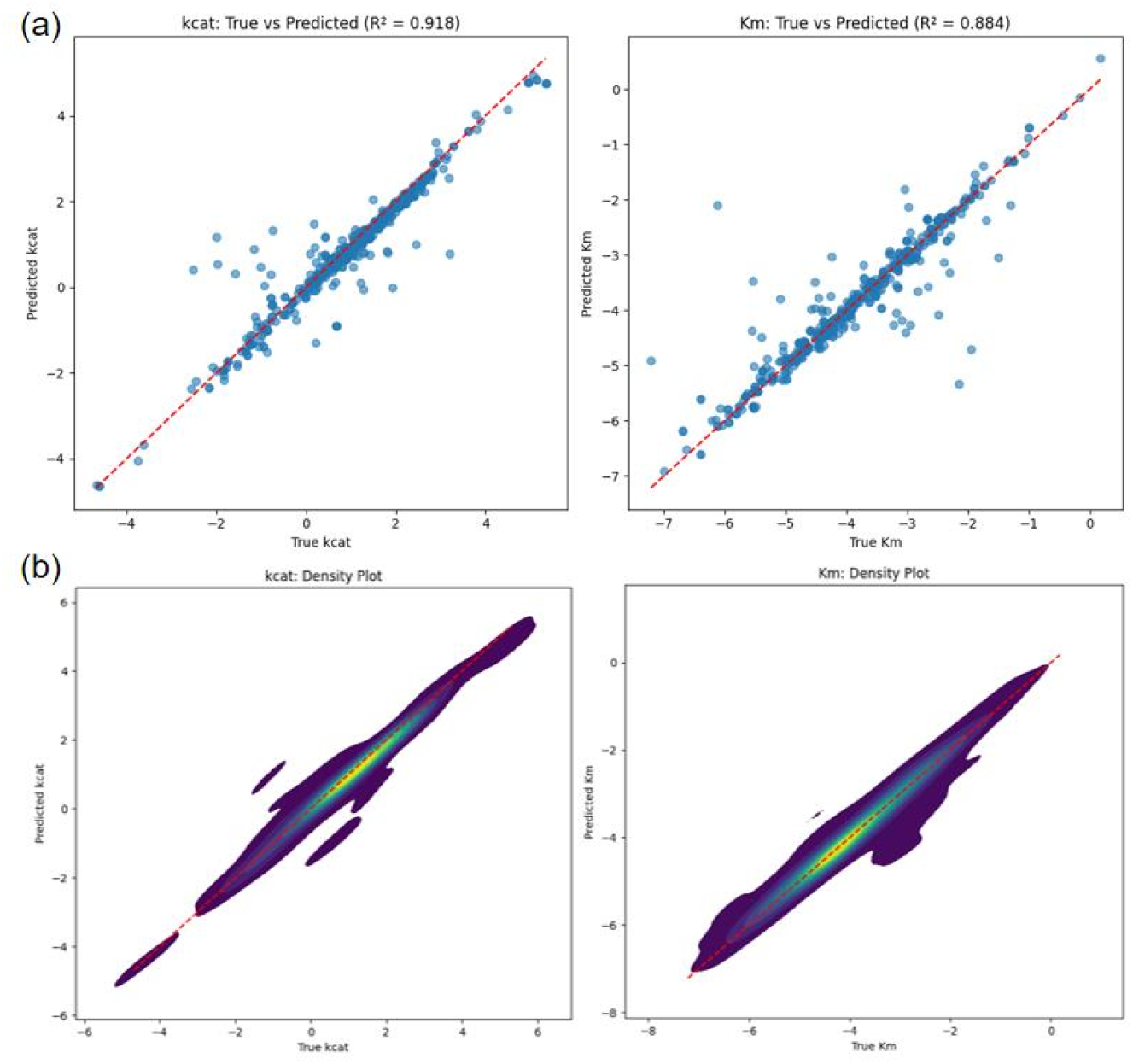
Scatter plots of true versus predicted values for (a) log_10_(*k*_*cat*_) and (b) log_10_(*K*_*M*_) on the random split validation set. The high correlation confirms the robustness of the multi-modal predictions.

1. **Structure Acquisition**: Enzyme structures are sourced from the PDB or predicted using ESMFold [7]. Substrate 3D conformers are generated from SMILES using RDKit with UFF energy minimization.
2. **Pocket Extraction**: We perform molecular docking using DiffDock [15], a diffusion-based docking method that automatically predicts binding poses without requiring manual specification of binding sites. The active pocket is defined as all protein atoms within 5.0 Å of the docked substrate.
3. **Graph Construction**: The pocket is converted into a radius graph (*r*_cut_ = 4.0 Å) for input into the GNN.

### 2.2 PocketGNN Architecture

PocketGNN employs a multi-branch architecture to fuse information across modalities.

#### 2.2.1 Local Branch: Geometry-Enhanced Graph Encoder

The active site is modeled as a geometric graph.

- **Node Features (52-dim)**: Detailed atom-level features including element type, residue identity, and electronic state descriptors, providing a rich chemical description of the micro-environment.
- **Geometric Edge Encoding (24-dim)**: To capture the stereochemistry, we employ a sophisticated edge featurization **e**_*ij*_ comprising:

- **Radial**: 16-dim Gaussian RBF encoding of distances.
- **Angular**: 4-dim features derived from bond angles (*θ*_*ijk*_) formed by neighbor pairs.
- **Torsional**: 4-dim features derived from dihedral angles, capturing the local flexibility and chirality.

A Graph Attention Network (GAT) [14] processes this graph, where attention weights are modulated by these geometric edge features, ensuring that the message passing respects the spatial configuration of the pocket.

#### 2.2.2 Global Branch: Sequence Semantics Fusion

To incorporate long-range dependencies, we utilize ESM-2 embeddings. For each enzyme, a global sequence vector **s**_seq_ is extracted. This vector encapsulates the evolutionary history and global structural propensities of the protein.

#### 2.2.3 Cross-Modal Late Fusion

The local structural representation **h**_pocket_ (from graph pooling) and the global sequence representation **s**_proj_ (from ESM-2 projection) are concatenated:

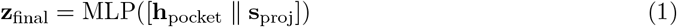

To avoid dimensional imbalance between the high-dimensional sequence embedding (640-1280 dim) and the graph representation (128-256 dim), we employ a projection layer that maps the ESM-2 embedding to the same dimension as the graph features before concatenation. This late fusion allows the model to dynamically weigh the immediate physical constraints of the pocket against the global stability implied by the sequence.

## 3 Experiments

### 3.1 Experimental Setup

The dataset is randomly split into 80% training and 20% validation sets for in-distribution evaluation. Additionally, to rigorously test generalization, we construct a homology-aware split using MMseqs2, ensuring that no sequence in the test set shares > 40% identity with the training set. We evaluate performance using Coefficient of Determination (*R*^2^), Pearson Correlation Coefficient (*r*), and Mean Squared Error (MSE).

### 3.2 Data Quality and Feature Analysis

To ensure that our graph representations capture meaningful physicochemical signals rather than spurious correlations, we performed a comprehensive data-level diagnosis. Pearson correlation analysis revealed that several geometric and electronic features exhibit significant correlation (| *r* | > 0.3) with catalytic rates. Furthermore, a Random Forest regressor trained solely on aggregated statistical descriptors achieved a test Pearson correlation of ~ 0.55. While this suggests that simple descriptors alone cannot fully solve the problem, it confirms the existence of *nonlinear but extractable* signal within the pocket geometry, validating our focus on the active site.

### 3.3 Performance Comparison

PocketGNN demonstrates superior performance, effectively leveraging its cross-modal design.

#### In-Distribution Performance (Random Split)

As shown in Table 1, the full model achieves a Pearson *r* of 0.98 and *R*^2^ of 0.918 for log_10_(*k*_*cat*_). The significant improvement over the “Structure-only” (PGNN-Basic) and “Sequence-only” baselines highlights the synergy of combining local geometry with global semantics. While an *R*^2^ > 0.9 is exceptionally high for kinetic data and may reflect the high similarity within enzyme families in a random split, the consistent ranking capability (*r* = 0.98) remains a strong indicator of model utility for in-family screening.

**Table 1:**
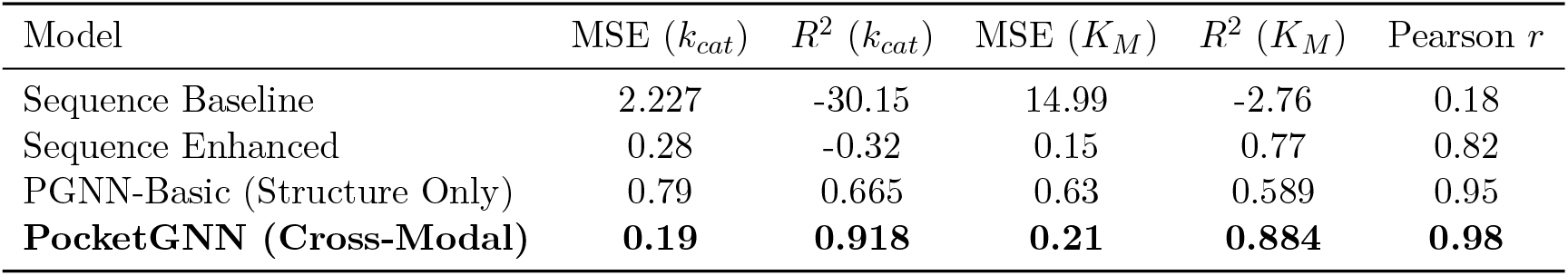
Quantitative prediction results on the random validation set.

#### Out-of-Distribution Generalization (Homology Split)

To assess robustness against data leakage, we evaluated the model on the strict 40% identity split. In this challenging zero-shot setting, PocketGNN maintains a Pearson correlation of **0.667** (with *R*^2^ = 0.437). While the *R*^2^ drop reflects the inherent difficulty of predicting kinetic rates for unseen enzyme families (OOD), the moderate-to-high correlation confirms that the model has learned generalizable physicochemical trends rather than merely memorizing sequence motifs.

#### Comparison with State-of-the-Art Methods

We compared PocketGNN against recent SOTA methods on the same 40% homology split test set (Table 2). PocketGNN achieves a Pearson correlation of 0.667, outperforming CatPred (0.52) and CataPro (0.497) by significant margins (28.7% and 34.2% relative improvement, respectively). This demonstrates the value of our pocket-centric geometric representation combined with cross-modal fusion, particularly in challenging OOD scenarios where sequence similarity is low.

**Table 2:**
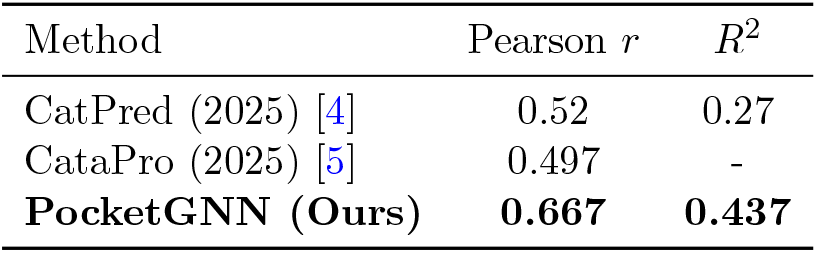
Comparison with SOTA methods on 40% homology OOD split.

| Method | Pearson $r$ | $R^2$ |
| --- | --- | --- |
| CatPred (2025) [4] | 0.52 | 0.27 |
| CataPro (2025) [5] | 0.497 | - |
| **PocketGNN (Ours)** | **0.667** | **0.437** |

Figure 2 visualizes the correlation between predicted and ground truth values, showing tight clustering along the diagonal.

### 3.4 Ablation Studies

- **Geometry Matters**: Removing the angle and dihedral features (reverting to simple distance-based GAT) resulted in a notable performance drop, confirming that “bag-of-distance” models are insufficient for capturing catalytic nuances.
- **Sequence Context Matters**: Removing the ESM-2 branch decreased accuracy, particularly for enzymes where the pocket structure alone might be ambiguous or flexible, proving the value of global evolutionary context.

### 3.5 EC-wise Performance Analysis

To further understand model behavior across enzyme classes, we analyzed performance metrics by EC number (Figure S1). The model achieves highest accuracy on Hydrolases (EC 3) and Transferases (EC 2), likely due to the abundance of structural data for these classes. Performance is slightly lower on Ligases (EC 6), reflecting the complexity of multi-substrate mechanisms. However, the consistent performance trend across major classes indicates that PocketGNN is not biased towards a single catalytic type.

### 3.6 Interpretability and Case Study

We validated the model’s focus using the Input × Gradient method on Human Aldehyde Dehydrogenase 2 (ALDH2, P30838). The visualization (Figure 3) shows that the model assigns high saliency (red) to the catalytic nucleophile (Cys302) and proton abstractor (Glu268), as well as the substrate binding cleft. Importantly, the high-saliency atoms are often connected by edges with strong angular and torsional feature weights, suggesting that PocketGNN leverages specific stereochemical constraints—not just proximity—to identify the chemical engine of the enzyme.

**Figure 3.**
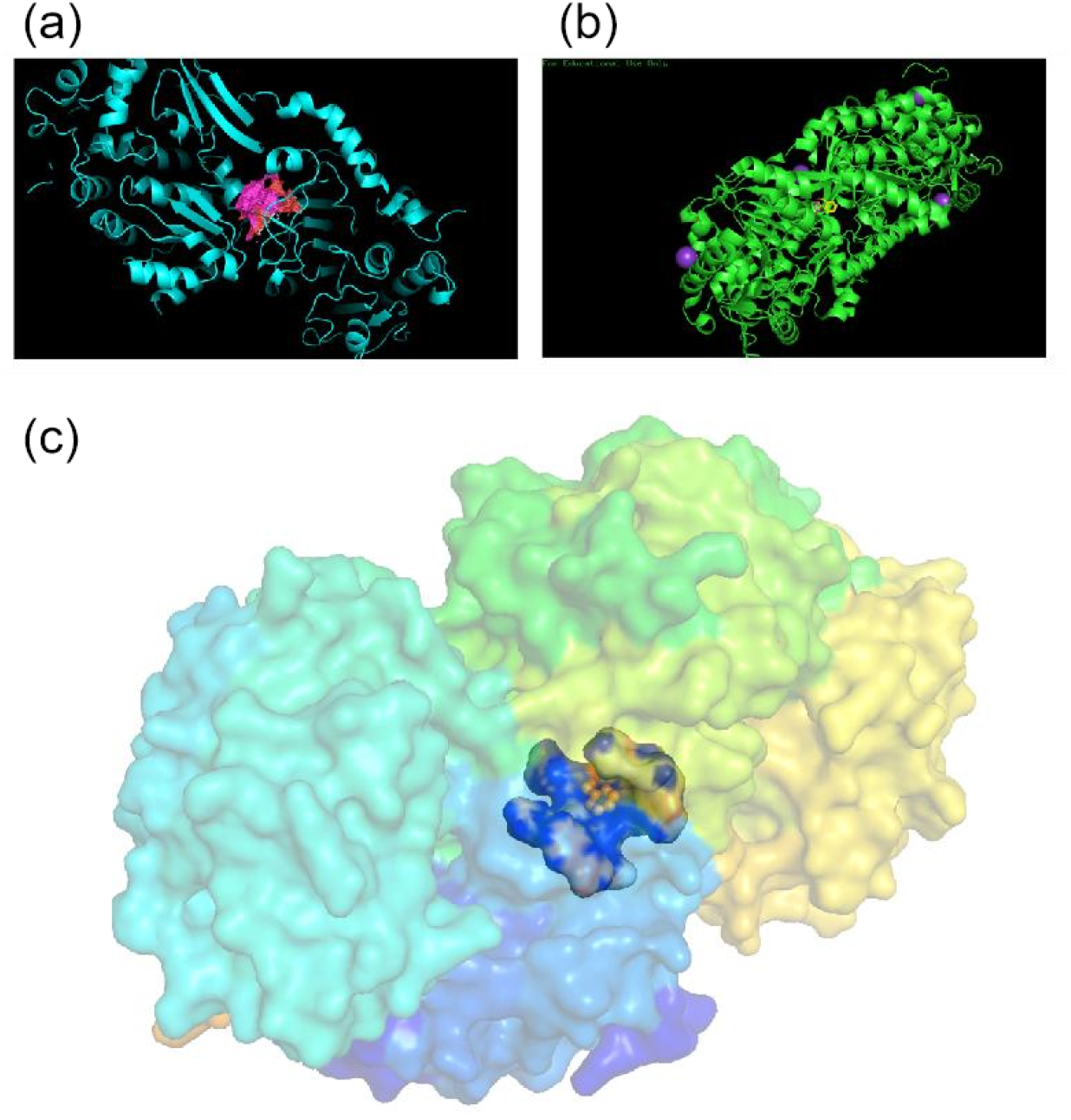
Visualization of atomic contributions for ALDH2. High-saliency atoms (red) coincide with key catalytic residues, demonstrating the model’s ability to identify the chemical engine of the enzyme.

### 3.7 Code and Data Availability

All code, trained models, and processed datasets are available at https://github.com/RektonLee/PGNN_final. The raw data is derived from IntEnzyDB [11], a publicly available database. Pretrained models and processed datasets are available upon reasonable request.

### 3.8 Limitations and Future Work

Our study has several limitations that warrant future investigation:

- **Experimental Conditions**: Unlike MPEK [16], we do not explicitly model temperature, pH, or organism-specific effects. This may limit accuracy when predicting kinetics under conditions different from training data.
- **Uncertainty Quantification**: Unlike CatPred [4], we do not provide prediction confidence intervals. Future work will incorporate quantile regression or Monte Carlo dropout for uncertainty estimation.
- **Multi-substrate Reactions**: Our current framework handles single-substrate reactions. Extending to multi-substrate systems (as in CatPred) would broaden applicability.
- **Docking Dependency**: The quality of pocket extraction depends on DiffDock’s accuracy. Future work will explore alternative docking methods (e.g., Chai-1) and pose validation strategies to improve robustness.
- **Data Quality**: Our study focuses on a curated high-quality dataset, and performance on low-quality or incomplete data remains to be tested.

While the homology split demonstrates generalization capability, the drop in *R*^2^ indicates room for improvement in transfer learning across disparate enzyme families. Future work could explore pre-training strategies specifically for pocket geometries to further bridge this gap.

## 4 Conclusion

PocketGNN represents a step forward in *cross-modal* enzyme modeling. By fusing the “short-range” precision of geometry-enhanced pocket graphs with the “long-range” breadth of protein language models, we achieve high-accuracy kinetic predictions (*r* = 0.98). Our rigorous evaluation under strict homology splitting further confirms that the model captures genuine physical signals (*r* = 0.67) beyond sequence memorization, outperforming recent SOTA methods by significant margins. This framework highlights the necessity of treating enzymes as both physical objects (3D) and evolutionary products (1D) to fully decode their catalytic potential.

